# Differences in the metabolite profiles of tender leaves of wheat, barley, rye and *Triticale* based on LC-MS

**DOI:** 10.1101/2020.12.03.409847

**Authors:** Piyi Xing, Zhenqiao Song, Xingfeng Li

## Abstract

Wheatgrass has emerged as a functional food source in recent years, but the detailed metabolomics basis for its health benefits remains poorly understood. In this study, liquid chromatography-mass spectrometry (LC-MS) analysis were used to study the metabolic profiling of seedlings from wheat, barley, rye and triticale, which revealed 1800 features in positive mode and 4303 features in negative mode. Principal component analysis (PCA) showed clear differences between species, and 164 differentially expressed metabolites (DEMs) were detected, including amino acids, organic acids, lipids, fatty acids, nucleic acids, flavonoids, amines, polyamines, vitamins, sugar derivatives and others. Unique metabolites in each species were identified. This study provides a glimpse into the metabolomics profiles of wheat and its wild relatives, which may form an important basis for nutrition, health and other parameters.

**Practical Application:** This manuscript present liquid chromatography-mass spectrometry (LC-MS) results of young sprouts of common wheat and its relatives. Our results may help to better understand the natural variation due to the genotype before metabolomics data are considered for application to wheatgrass and can provide a basis (assessment) for its potential pharmaceutical and nutritional value.

## Introduction

*Triticum aestivum* L. is a species within a genus of annual and biennial grasses that is widely cultivated throughout the world as an important grain crop. In addition to its seed, wheatgrass as functional food has gained popularity all over the world (Singh et al. 2012; Anwar et al., 2015; Akbas et al., 2017; Parit et al., 2018). Wheatgrass refers to the young grass of the common wheat plant obtained following germination of the wheat seed for 6–10 days and generally consists of a greatly compressed stem or crown and numerous narrowly linear or linear-lanceolate leaves.

A survey of 1000 plants for their pharmacological and therapeutic properties in 2010 revealed that wheatgrass has pharmacological significance (Cravotto et al., 2010). In previous literature, wheatgrass was shown to have anti-aging, antioxidant, healing and blood-clotting properties (Singh et al., 2012); an aqueous extract of wheatgrass was even shown to have anti-cancer properties (Arora and Tandon, 2016). Wheatgrass is becoming one of the most widely used health foods. At present, wheatgrass is available in various forms (fresh juice, frozen juice, tablets, and powders) as a part of a healthy diet in the USA, East Asian countries and Eastern Europe (Parit et al., 2018). Apart from wheatgrass, barley grass (BG), considered a useful food for humans, has recently been receiving considerable attention for its ability to help prevent various diseases (Lahouar et al., 2015).

The use of sprouts and young wheat plantlets in human nutrition is increasing because these materials often contain phytochemicals and other high-value nutrients. In previous studies, wheatgrass was found to contain abundant chlorophyll; the minerals K, Ca, Fe, Mg, Na and S; vitamins such as A, B, C and E; enzymes and 17 amino acids (Singh et al., 2012). Wheatgrass is a rich source of tocopherols with high vitamin E potency. Phytochemical screening and gas chromatography-mass spectrometry (GC-MS) analysis in aqueous extracts of wheatgrass has verified the presence of bioactive components, such as flavonoids, triterpenoids, anthraquinol, alkaloids, tannins, saponins, sterols, squalene, caryophyllene and amyrins, many of which have demonstrated antioxidant activity (Kothari et al., 2011; Durairaj et al., 2014). These previous studies focused on aqueous wheatgrass extracts, but comprehensive metabolomics data are limited for the methanol extract. Moreover, these previous studies of the bioactive metabolites in wheatgrass mainly used a targeted approach to detect the presence or concentration of known compounds, which limits the detection to a relatively small portion of the metabolites. Therefore, the quantity and detailed composition of metabolites in extracts of wheatgrass remains unclear, and information about the broad metabolomics basis for the health benefits of wheatgrass has been limited to date.

Metabolomics technology offers the capability to identify and measure hundreds of separate biochemical entities simultaneously. For this reason, the adoption of metabolomics to study the nutrient content of grains has been recently gaining popularity (Lee et al., 2013). Untargeted metabolomics provides a base from which to discover the mechanisms behind pharmacological effects and to reveal the important basis for grasses as a food supply. Comprehensive metabolomic profiling requires a reliable protocol that enables the extraction of maximum metabolic information in a high-throughput and reproducible manner to cover as broad range of metabolites as possible. Untargeted metabolomics analyses are performed with a variety of analytical methods including GC-MS (Tarpley et al., 2005), nuclear magnetic resonance (NMR) spectroscopy (Heuberger et al., 2014) and liquid chromatography-mass spectrometry (LC-MS) (Metz et al., 2007; De Vos et al., 2007). LC-MS has several advantages, including greater sensitivity and dynamic range; does not require sample derivatization; and provides analytical access to molecules of more diverse mass and chemical structures, such as secondary metabolites.

Most previous studies performed a component analysis of only wheat or barley. Other important cereal grains such as rye and triticale have demonstrated functions in human health, but few reports of metabolic research on their grasses exist (ÖZKÖSE et al., 2016). Moreover, a comparison of different cereals has not been reported to date. Further studies are needed to better understand how classes of phytochemicals differ between cereal species and which classes contribute to the maintenance of human health.

In the present study, an untargeted LC-MS-based metabolomics approach was applied to investigate the grasses of barley, wheat, rye and Triticale grown in China. The aim was to search for discriminative metabolites in wheatgrass, with an emphasis on the differences among wheat, rye and durum, to gain further insights into their possible roles in the beneficial health effects. Our results may help to better understand the natural variation due to the genotype before metabolomics data are considered for application to wheatgrass and can provide a basis (assessment) for its potential pharmaceutical and nutritional value.

## Materials and Methods

### Plant materials

Four species were used: *Secale cereal*, octoploid *Triticale, Hordeum vulgare*, and *T. aestivum*, including three cultivars, ‘Huixianhong’ (HXH), ‘Yannong 15’ (YN), ‘Jimai 22’ (JM). The material code names of *S. cereal*, octoploid *Triticale*, and *H. vulgare*, were represented as 109, 221 and BAR, respectively.

### Seedling germination

The grass seeds were sown, germinated and raised in pots under greenhouse conditions. The experiment was set up as a completely randomized design with eight replications. The plants were grown without fertilizer. Plastic pots of 22 × 20 cm were filled with 5 L of sterilized peat to 1 cm below the pot brim. Throughout the experiment, the plants were irrigated every day with deionized water. The wheatgrass was cut on the 10th day. The freshly cut grass (200 g) from each cultivar with a height of 10 cm was collected, immediately frozen in liquid nitrogen, ground into power, and stored at −80 °C for further analysis.

### Extraction and analysis of metabolites

All chemicals used were of analytical grade. Methanol and formic acid from Merck (Darmstadt, Germany) were used for extraction and separation, respectively. The ultra gradient-grade acetonitrile and methanol used in the chromatographic experiments were from Romil (Cambridge, UK). Deionized water was prepared by a Milli-Q system from Millipore (Bedford, MA, USA).

For LC-MS analysis, 50 mg of a tissue sample was transferred to a 1.5-mL centrifuge tube and submerged in 0.8 mL of methanol (Darmstadt, Germany), followed by the addition of 100 µL of L-2-chlorophenylalanine as an internal standard. The mixture was then crushed for 60 s at 60 Hz using a tissue crushing apparatus, followed by scrolling for 30 s, and then centrifugation at 12,000 rpm for 15 min on 4 °C. Next, 200 µL of the supernatant was collected from each sample into a vial (4 mL) for further analysis. Eight biological replicates were prepared for each genotype, resulting in a total of 40 extracts. Quality control samples were prepared by pooling the material of several randomly selected samples to assess the technical variation, with 8 replicates. A volume of 4 µL was injected into the LC-MS instrument for analysis. The quality control samples were injected after every 10 sample extracts.

LC-MS (Thermo, Ultimate 3000 LC, Orbitrap Elite) was used for metabolite profiling (instrument analysis platform in this experiment). The sample components were separated on a C18 olumn (Hypersil GOLD C18; 4.6 mm × 100 mm; 3 µm particle size; Agilent, Santa Cruz, CA, USA). The column was eluted at 0.3 mL/min with a gradient from 95% solvent A (0.1% formic acid in water) and 5% solvent B (0.1% formic acid in acetonitrile) for the first 2 min, followed by a gradual shift to 5% solvent A and 95% solvent B by the 15th min and to 95% solvent A and 5% solvent B by the 17th min. The column temperature was maintained at 40 °C. The injection volume was set at 4 µL for each sample.

The optimized electrospray ionization parameters were set as follows: heater temperature of 300 °C, sheath gas flow rate of 45 arb, auxiliary gas flow rate of 15 arb, sweep gas flow rate of 1 arb, spray voltage of 3.0 kV (positive mode) or 3.2 kV (negative mode), capillary temp of 350 °C, and S-lens RF level of 30% (positive mode) or 60% (negative mode). The spectral mass range was 50–900 (*m*/*z*).

### Multivariate data analysis

The mass, retention time, and abundance of metabolic features were obtained. The resulting LC-MS data were first processed by normalizing the peak area of each analyte based on the total integral area calculation using an in-house script (Microsoft Office Excel). All processed data were submitted to principal component analysis (PCA) to identify clustering trends. QC samples were analyzed at constant intervals to ensure that the data acquisition for the LC-MS metabolic profiling was reproducible for all samples. The relative content of a compound was determined according to its normalized peak area.

PCA and orthogonal projection to latent structure with discriminant analysis (OPLS-DA) were carried out using SIMCA-P + 13.0.2 software (Umetrics, Sweden) (Dai et al., 2010). PCA was performed with mean center scaling, and the data are presented in the form of a scores plot. OPLS-DA results were obtained with unit variance scaling while regarding the class information as Y variables and are shown in the form of a scores plot. The quality of the model was estimated from the parameters of the cross-validation: R2X, which shows the total explained variables for all LC-MS data, and Q2, which shows the predictability of the model.

The OPLS-DA model VIP (variable importance in projection) value (threshold >1) was combined with the t-test value of P (P < 0.05) to identify differentially expressed metabolites (DEMs). The qualitative method to identify DEMs involves searching online databases (http://metlin.scripps.edu/) based on *m*/*z* or accurate molecular mass information. Hierarchical cluster analysis (HCA) was performed to determine the metabolite expression in different species (Saeed et al., 2003). Figure 2 presents the relationship among metabolites based on HCA with data obtained in the positive and negative ionization modes and the relationship among species.

## Results

### PCA of metabolic profiles

An untargeted evaluation strategy was implemented for all 48 tissue samples (8 samples per cultivar, six cultivars). Eight QC samples for LC analysis were inserted into the analytical sequence of real samples, and the QC chromatograms (Supplemental figures S1 and S2) reflect the reliability of the instrumental analysis. The total ion chromatograms of the metabolomic analysis of HXH are shown in Supplemental figures S3 and S4. Based on the retention times and *m*/*z* values, 1810 feature peaks in positive ionization mode and 4403 features in negative ionization mode were observed. PCA can be used to elucidate the differences among samples, as shown in Figures 1A and 1B. The QC samples concentrated in the center of the coordinate axes, reflecting the reliability of LC-MS.

**Figure 1.**
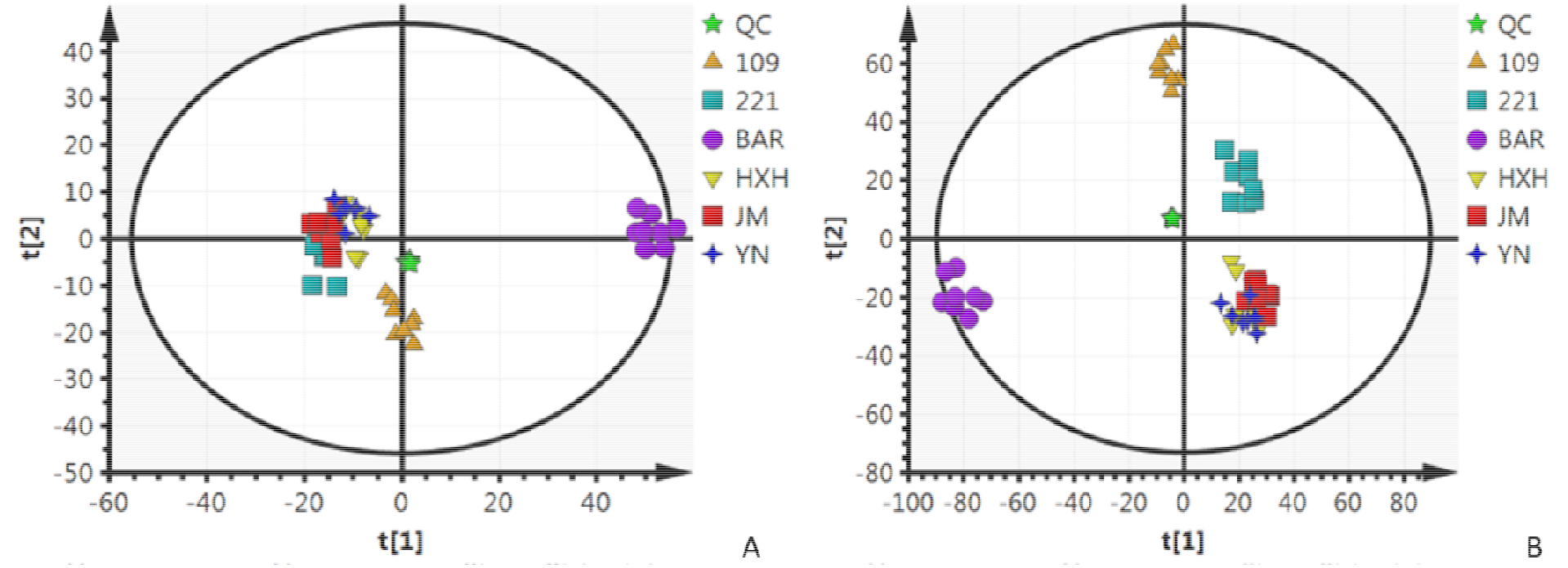
PCA of the metabolic profiles of four cereal grasses. Note, A. positive mode; B. negative mode; Each symbol represents one chromatogram. Vectors are loadings.

**Figure 2.**
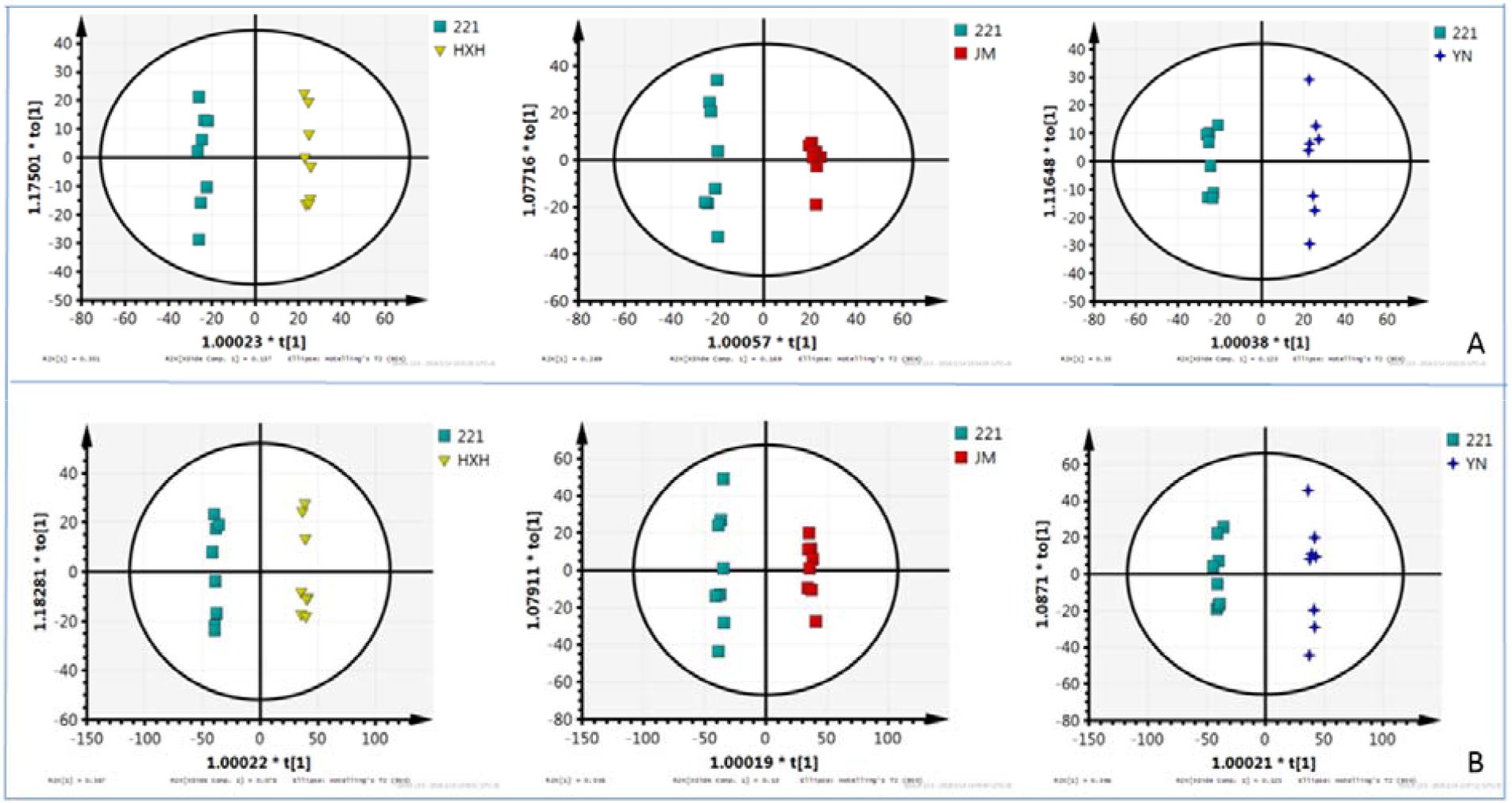
OPLS-PCA between triticale (221) and the three wheat cultivars (HXH, JM, and YN). Note, A. positive mode; B. negative mode.

The PCA representation displays the LC-MS/MS base peak chromatograms in the first and second principal component coordinates. In positive mode, 10 principal components were obtained with cumulative R2X and Q2 values of 0.846 and 0.616, respectively. For the unsupervised model analysis, the main quality parameter of the discriminant model, R2X, is generally greater than 0.4, indicating that the model is reliable. The first and second principal components explained 26.3% and 18.1% of the variation, which separated barley (BAR) and rye from the wheat cultivars, but triticale and the three wheat cultivars greatly overlapped (Fig. 1A). These results illustrate that the metabolites of barley, rye, and wheat were significantly different in positive mode but the differences between triticale and the wheat cultivars were not obvious. In negative mode (Fig. 1B), 9 principal components were obtained with cumulative R2X and Q2 values of 0.786 and 0.645, respectively. PC1 and PC2 accounted for 28.4% and 18.9% of the variation, respectively, illustrating a distinction between barley, rye, triticale and the three wheat cultivars; in particular, triticale can be clearly visually separated from rye and wheat. These results demonstrate the better ability of negative mode to differentiate among the samples compared to positive mode.

### OPLS-PCA of metabolism

OPLS-DA was used to highlight the separation of two special groups and demonstrated an enormous separation of different species along the t[1] axis (figure 2, supplemental figures S5 and S6). Figure 2 shows the OPLS-PCA of metabolite data from triticale and three wheat cultivars. In positive mode, the first principal component explained 48.9%, 45.7% and 47.3% of the variation separating triticale from the three wheat cultivars HXH, JM and YN, respectively (Fig. 2A). In negative mode, the first principal component explained 44.5%, 46.6% and 52.1% of the respective variation (Fig. 2B). Similar results were observed between rye and the three wheat cultivars and between barley and the three wheat cultivars (Supplemental figures S5 and S6).

### Metabolites with different expression levels distinguished between two groups

Using the VIP and t-test values in the OPLS-DA model, 109 metabolites in positive mode and 124 metabolites in negative mode annotated DEMs between these species were selected, 69 of which were simultaneously detected under the two modes (Table 1). The majority of compounds were shared across at least two grass genotypes, and statistical analysis showed that the differences were significant between the genotypes. The detailed metabolites are listed in supplemental tables S1 and S2.

**Table 1.**
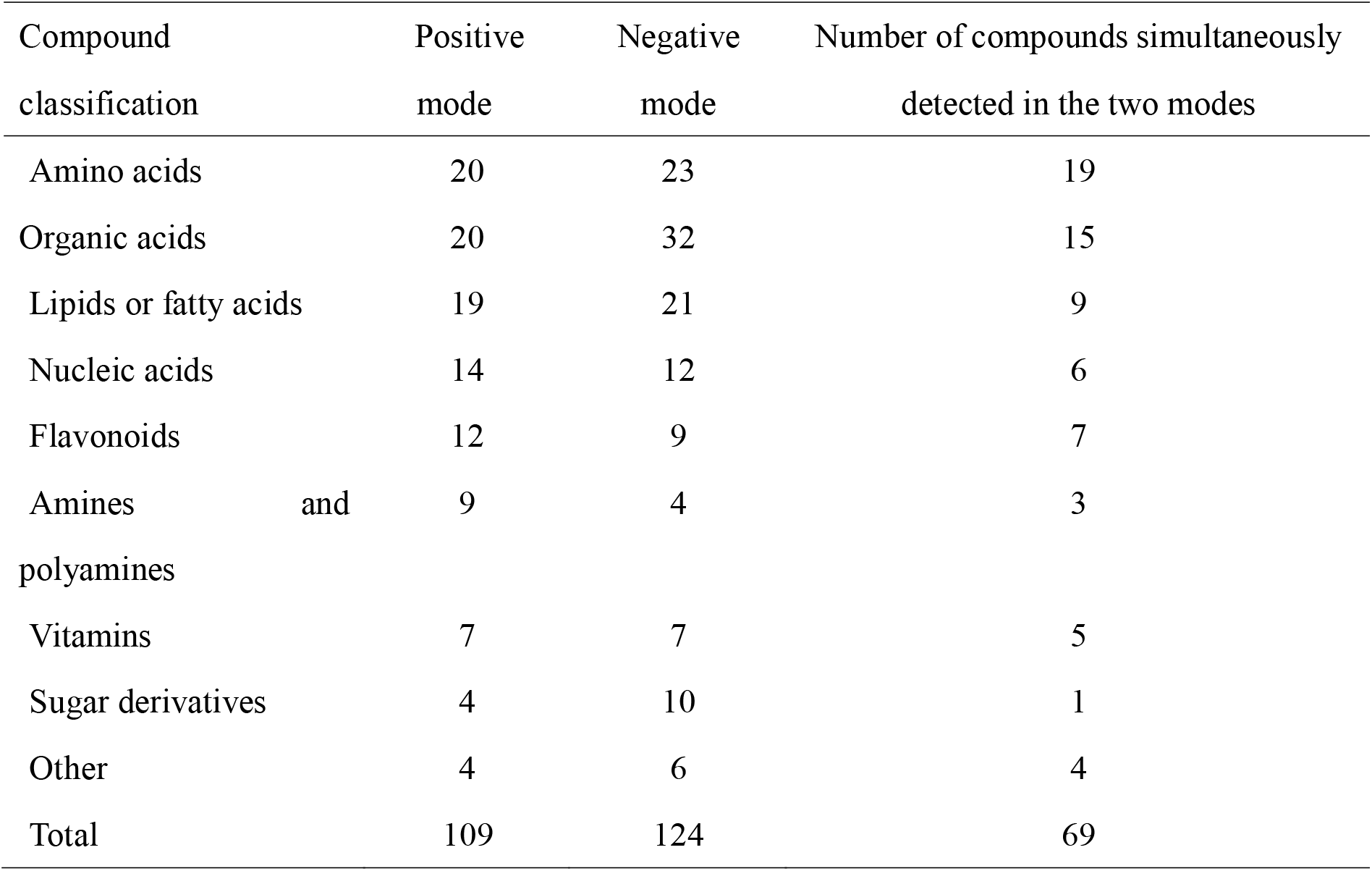
DEM classification in metabolic profiling.

The classification scheme illustrates the distribution, number and diversity of compound classes detected in the various species. Twenty-two amino acids were detected in all four species, including eight essential amino acids, with valine, phenylalanine, lysine and isoleucine at relatively high levels, consistent with a previous report (Anwar et al., 2015). Two semi-essential amino acids (histidine and arginine) and four non-essential amino acids (L-threonine, L-glutamate, L-aspartic acid, and proline) were also observed. Seven vitamins were detected in the four species, mainly including C and B vitamins such as riboflavin (vitamin B2), niacin (vitamin B3), pantothenic acid (vitamin B5), pyridoxal (vitamin B6), D-biotin (vitamin B7) and dehydroascorbic acid (vitamin C). Vitamin C was detected at high levels in all species. In addition, we found 12 flavonoid compounds, including catechin, coumarin, kaempferol, pheophorbide a, quercitrin, rutin, quercitrin, and baicalin. Coumarin and catechin had relatively high expression levels in four species, indicating that aqueous extracts of cereal grasses are good sources of antioxidants.

Previous studies reported few polyphenols and flavonoids, such as reduced ascorbic acid, quercetin, rutin, reduced glutathione or ferulic acid and vanillic acid, in wheatgrass (Calzuola et al., 2004; Yang et al., 2001). In the present paper, more than 20 organic acids were detected, with high amounts of ferulic acid and malic acid, and 15 were simultaneously detected both in positive and negative mode. Five organic acids were only detected in positive mode, with high expression of γ-aminobutyric acid, but 16 were detected in negative mode, with high expression of quinic acid, glutaconic acid and pyruvate in four species. The metabolomics data also included 14 nucleotides, 20 fatty acids or lipids, and a few amines or polyamines, as well as a small proportion of other compounds. The LC-MS analysis confirmed the presence of primary metabolites, such as monosaccharides or monosaccharide derivatives such as nucleotides, vitamins, amino acids, and fatty acids; a variety of lipids and other vital substances; and diverse secondary metabolites. The primary metabolites were more abundant than the secondary metabolites such as flavonoids, which indicates that grasses may possess more primary metabolic pathways.

### Cluster analysis of the DEMs in positive mode

Based on the relative contents of metabolites, HCA was applied to display the distribution of the DEMs across genotypes, as shown in Figure 3A. Four clusters were identified with distinct patterns of altered metabolite abundances in Figure 3A in positive mode.

**Figure 3.**
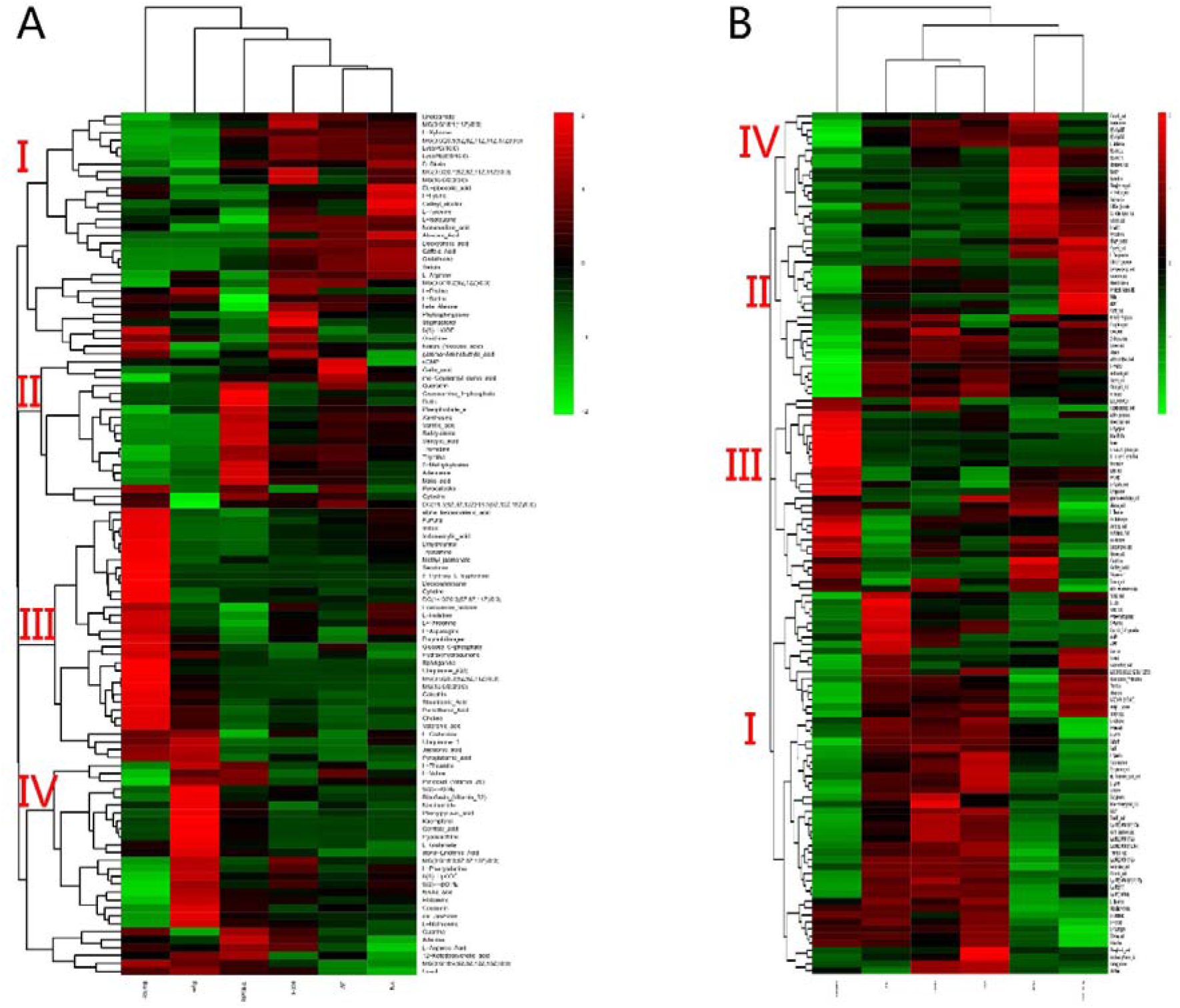
HCA clusters of the metabolites across genotypes. Note, A, positive mode; B, negative mode.

Group I included 23 compounds, including five amino acids, six kinds of organic acids and two kinds of flavonoids, that obviously had higher expression in the three wheat cultivars than in the other species (supplemental table S1). Some of these compounds, including gallic acid and p-coumaroyl quinic acid, were highly co-expressed in wheat and triticale (supplemental table S1), and polyamines (phytosphingosine) were more abundant in wheat and barley (supplemental table S1).

Group II consisted of 19 compounds, which had higher expression in triticale and included three flavonoids (quercitrin, rutin, and pheophorbide a), three organic acids (vanillic acid, salicylic acid, and malic acid), six nucleic acids, salicylamide, glucosamine-1-phosphate, and pyrocatechol.

The third group comprised approximately 40 compounds with higher contents in barley, including six amino acids, six organic acids, seven lipids or fatty acids, four nucleic acids, four vitamins and four amines or polyamines. Methyl jasmonate and catechin had higher contents in BG. L-asparagine, indole and choline were detected in relatively high amounts.

Approximately 20 compounds, including five amino acids, five fatty acids, two organic acids (ferulic acid and gentisic acid), pyridoxal (vitamin B6), riboflavin (vitamin B2), histamine and niacinamide, were present at high levels in rye (supplemental table S1) and clustered into the fourth group. 12-Ketodeoxycholic acid and cis-jasmone had higher contents in both rye and triticale (supplemental table S1),

In positive mode, some metabolites were highly expressed in all the species, including some amino acids (L-isoleucine, L-valine, and L-phenylalanine), fatty acids (indoleacrylic acid, 9(S)-HOTrE, 9(S)-HpOTrE, and 9(S)-HpODE), nucleic acids (adenine, thymine, and guanine) and organic acids (malic acid, ferulic acid and γ-aminobutyric acid). These distributions of DEMs show that the main compound forms were different in the different species.

### Cluster analysis of the DEMs in negative mode

As seen in Figure 3B, which is based on the data obtained in negative mode, the compounds clustered into 4 groups.

The first group included approximately 54 compounds, of which 30 had relatively high expression in the three wheat cultivars (supplemental table S2), including seven amino acids, six organic acids, eight fatty acids or lipids, four nucleic acids, one vitamin (nicotinic acid) and one flavonoid (catechin).

Some of the compounds in the first group were highly co-expressed in wheat and triticale (supplemental table S2), including acetyl-L-tyrosine, five organic acids (acetoacetic acid, ferulic acid, chlorogenic acid, salicylic acid, and sinapic acid), five fatty acids (oleic acid, α-linolenic acid, MG(0:0/18:1(11Z)/0:0), linoleic acid, LysoPE(0:0/20:1(11Z)), and MG(0:0/22:5(4Z,7Z,10Z,13Z,16Z)/0:0)), three nucleic acids and two sugars (sedoheptulose-7-phosphate and L-xylulose).

Three amino acids (L-glutamine, L-histidine, and L-asparagine) and two polyamines (histamine and phosphorylcholine) had higher contents in wheat and barley (supplemental table S2).

Among the three wheat cultivars, JM22 contained unique compounds with higher contents compared with the other cultivars and species (in red and italics in table 4): L-glutamate, vanillic acid, gallic acid, MG(0:0/15:0/0:0), cAMP, cGMP, quercitrin, D-mannitol, quercetin-3-O-glucoside, hydroxyhydroquinone, and L-dopa. This result illuminates the wide range of germplasm diversity in cereal grasses.

The following 27 compounds were expressed much more highly in barley (supplemental table S2): eight amino acids (L-tryptophan, β-alanine, N-acetyl-L-glutamic acid, 5-hydroxy-L-tryptophan, ketoleucine, L-aspartic acid, L-theanine, and oxidized glutathione), 12 organic acids (octadecanedioic acid, methyl jasmonate, indoleacrylic acid, malic acid, pyruvate, γ-aminobutyric acid, abscisic acid, jasmonic acid, dodecanedioic acid, succinic acid, fumaric acid, and α-ketoisovaleric acid), two vitamins (pantothenic acid and ubiquinone-1), two flavonoids (cis-jasmone and caffeyl alcohol), levoglucosan, indole and porphobilinogen.

Only approximately 10 compounds in triticale were detected at high levels based on their normalized peak areas (supplemental table S2): two organic acids (quinic acid and glutaconic acid), dGMP, two vitamins (dehydroascorbic acid and pyridoxal (vitamin B6)), two flavonoids (rutin and pheophorbide a), and two sugars (glucoheptonic acid and DIMBOA glucoside).

In rye, approximately 10 compounds were present at relatively high levels (supplemental table S2): L-methionine, five fatty acids or lipids (9(S)-HpODE, 9(S)-HOTrE, 9(S)-HODE, 9(S)-HpOTrE, and stearidonic acid), baicalin, 4-pyridoxic acid, gluconic acid, sinapyl alcohol, and nicotianamine. Approximately 10 compounds had higher contents in both rye and triticale (supplemental table S2), such as L-valine, L-phenylalanine, flavonoids (kaempferol), four organic acids (phenylpyruvic acid, DL-α-lipoic acid, gentisic acid and glyoxylic acid), DIBOA glucoside, salicylamide, and pyrocatechol.

In negative mode, several compounds were present in high amounts in all the species, such as L-isoleucine, L-tryptophan, nonanedioic acid, octadecanedioic acid, malic acid, quinic acid, glutaconic acid, oxoglutaric acid, 9(S)-HpODE, 9(S)-HOTrE, 9(S)-HODE, 9(S)-HpOTrE, stearidonic acid, and thymidine, which are mainly involved in primary metabolic pathways.

In both positive mode and negative mode, L-isoleucine, 9(S)-HOTrE, 9(S)-HpOTrE, and malic acid were simultaneously detected at high levels in all grass species, which may indicate that these are important substrates for many biochemical pathways.

### Genetic relationships among species

Figure 3A, 3B also presents the relationship among species based on the metabolomics data obtained under positive mode and negative mode. The two maps show similar clusters for the different genotypes. The three wheat cultivars clustered first, which clearly indicates that they have similar metabolic expression characters, consistent with the results of Figure 1 in which the three wheat cultivars were hardly separated. Next, triticale was clustered with wheat, or triticale and rye were clustered into one group. The last species to cluster was barley, which reflects a more distant relationship between barley and the other species. The clustering results in Figure 3A, 3B further confirm the relationship among species presented by Figure 1 and is consistent with the genetic relationship based on chromosome composition. Therefore, the clustering analysis based on chemical composition is feasible and credible.

## Discussion

### LC-MS for untargeted and quantitative analyses of cereal grasses

Metabolomic approaches have emerged as a valuable tool for the plant sciences, especially for the discovery and identification of markers of characterization of different genotypes/phenotypes. In the past few years, the rapid development of high-throughput tools for metabolic profiling, such as detection of the levels of multiple metabolites in a single extract, has facilitated the analysis of a broad range of metabolites. Grass metabolomics is a promising approach to reveal the biochemical and genetic backgrounds of quality traits and may open a new door towards targeted functional food development. Using LC-MS for untargeted and quantitative analyses has gained considerable attention within the metabolomics community (Fernie et al., 2008; Cevallos-Cevallos et al., 2009). This study applied LC-MS to the profiling of a broad range of metabolites in grasses of wheat, barley, rye and triticale. The distinct differences in metabolite composition were highlighted among the tested genotypes. This research contributes to the characterization of species through the comparative profiling of metabolites found in different varieties, and metabolomics offers insight into grass metabolic fluctuations in cereal cultivars.

### Metabolite composition of cereal grasses

In this study, grasses from 6 wheat cultivars were used to explore the metabolite profile. We found abundant metabolites in these grasses: 1810 feature peaks in positive ionization mode and 4403 features in negative mode. These data will provide the material basis for further development of functional foods.

Germination and sprouting cause plants to synthesize metabolites such as vitamins, antioxidants and phenolics (Kulkarni et al., 2006). The identified metabolites mainly consisted of 20-23 amino acids, 20-32 organic acids, 20 fatty acids or lipids, 10 flavonoids, and 7 vitamins, as well as a few amines, polyamines and other compounds. The constituents were generally consistent with the report of Durairaj et al. (2014). The LC-MS analysis confirmed the presence of primary metabolites such as monosaccharides and monosaccharide derivatives including nucleotides, vitamins, amino acids, and fatty acids, as well as a variety of lipids, other vital substances and diverse secondary metabolites. Most of these compounds, such as amino acids, organic acids, bioflavonoids, and plant hormones, are bioactive and reported to be important or beneficial to human health. Young leaves and sprouts have increased nutritional value resulting from the natural sprouting of cereal seeds.

Amino acids are necessary for the continuous cell building, cell regeneration, and energy production needed for good health (Foda, 2012). In the present work, 22 amino acids were detected in all four species, with valine, phenylalanine, lysine and isoleucine present at high levels. These results are consistent with a previous study (Anwar et al., 2015) reporting 17 amino acids in wheatgrass juice. Chavan and Kadam (1989) stated that an increase in proteolytic activity during sprouting is desirable for nutritional improvement of cereals because it leads to hydrolysis of prolamins and the liberated amino acids such as glutamic and proline are converted to limiting amino acids such as lysine.

In the present study, more than 20 organic acids were detected. The previous studies on wheatgrass mainly detected few polyphenols and flavonoids, such as reduced ascorbic acid, quercetin, rutin, reduced glutathione, ferulic acid and vanillic acid (Calzuola et al., 2004). The present work is the first to detect such a high number of organic acids. Among the detected organic acids, ferulic acid and malic acid were highly abundant in all species, consistent with a previous study on cereal grain (Khakimov et al., 2014). During germination, some biologically active compounds such as ferulic and vanillic acids are synthesized in wheat sprouts, with concentrations reaching their maxima after 7 d. An increase in free phenolic acids with sprouting was also observed by Van Hung et al. (2011) and is supported by findings in rice, barley, sorghum, rye and oat (Donkor et al., 2012; Benincasa et al., 2015).

Twelve flavonoid compounds were detected, and coumarin and catechin were relatively highly expressed in the four species.

The four species grasses mainly contained C and B vitamins, consistent with previous reports on wheatgrass (Halime et al., 2015) and barley (Edwin and Sheeja., 2006). Vitamin C, as part of the secondary defense system, has scavenging properties against an overproduction of free radicals that may incidentally occur during a cell’s life cycle. B vitamins promote the metabolism in the body to convert sugar, fat, protein and other substances into heat when necessary.

Among the metabolites, α-linolenic acid, 9(S)-HpODE, and thymidine were expressed in high amounts in all the species. Many studies have shown that water extracts of wheatgrass are a good source of antioxidants. Wheatgrass extracts can be used as a dietary supplement for antioxidant compounds such as vitamins, polyphenols and flavonoids (Singh., 2017). Based on the obtained results, we conclude that aqueous wheatgrass extract contains various effective compounds and is a potential source of natural antioxidants. Further analysis of these species will uncover new effective compounds of pharmacological significance.

### Diversity of the metabolite compositions of different species

The phytochemical screening revealed variations in the phytochemical content among the grass species, which will lay the foundation for the development of diverse health foods. Only a small portion of the metabolites were not detected in one species, for example, baicalin, DIMBOA-glucoside, MG(0:0/18:1(11Z)/0:0), hypoxanthine, deoxycholic acid, quercitrin and rutin were not detected in BG. Most of the metabolites were detected in all species, showing that these cereal grasses had similar metabolic characteristics but most metabolites had significantly different expression levels.

For the grasses of the three wheat cultivars, the metabolomics data revealed such similar expression patterns that the cultivars first clustered into one group in Figure 2A, B. Among the 109 DEMs obtained in positive mode and 124 in negative mode, wheatgrass had 28 and 60, accounting for 25.68% and 48.39%, respectively, with higher expression compared with the other three species. These metabolites mainly consisted of amino acids, organic acids, nucleic acids and lipids or fatty acids, including essential nutrients such as L-lysine and important active chemical constituents such as nicotinic acid and catechin. Ethanol extracts of wheatgrass were found to have higher phenolic and flavonoid contents than aqueous extracts (Kulkarni et al., 2006). DIBOA-glucoside, MG(0:0/18:1(11Z)/0:0), and DIMBOA-glucoside, as well as high levels of deoxycholic acid and stearidonic acid, were detected in wheat. Catechins impart a puckering astringency and roughness to the oral cavity, and their effective antioxidant activity results from their special structures containing vicinal dihydroxy or trihydroxy groups. Considering its rich phenolic metabolites, the addition of 7.5% wheatgrass powder in muffins as a supplement provided beneficial health effects (Rahman et al., 2015).

In barley, approximately 7 compounds were not detected, while 40 compounds in positive mode and 27 in negative mode, accounting for 36.70% and 21.77%, respectively, had higher contents than in the other species. These metabolites primarily consisted of amino acids, organic acids, nucleic acids and lipids or fatty acid. Some vitamins such as pantothenic acid and ubiquinone-1 had higher expression in BG. Previous studies reported that BG contains a wide spectrum of vitamins as well as eight essential amino acids (Lahouar et al., 2015). Bioactive metabolites such as p-coumaroyl quinic acid, baicalin and kaempferol had higher expression, and kaempferol has antibacterial, anti-inflammatory, antitussive and expectorant effects. Because it possesses pharmacological activities such as anticancer, antioxidant and anti-inflammatory activity, barley grass powder has recently seen an upswing in its worldwide popularity and is continually being added the multitude of health products, such as green supplements and smoothies (Foda., 2012). In the USA, green barley is on the market as a food supplement.

In rye, approximately 20 compounds were detected in higher amounts in positive mode, including amino acids (L-valine, L-threonine, L-glutamate, L-methionine, and L-phenylalanine), fatty acids (α-linolenic acid, 9(S)-HOTrE, 9(S)-HpOTrE, 9(S)-HpODE, and MG(0:0/18:3(6Z,9Z,12Z)/0:0)), organic acids (such as ferulic acid and gentisic acid), and riboflavin (vitamin B2). In negative mode, only 10 compounds were detected in higher amounts. In both positive and negative mode, L-methionine, 9(S)-HpODE, 9(S)-HOTrE, 9(S)-HpOTrE, and pyridoxal (vitamin B6) were detected at higher levels in rye, levoglucosan and dodecanedioic acid were especially enriched in rye.

In triticale, approximately 15 metabolites in positive mode and 10 in negative mode were detected at high levels, including organic acids, vitamins, and flavonoids. Notably, two flavonoids, rutin and pheophorbide, were particularly enriched in triticale. Rutin can help combat inflammation-related skin damage. Therefore, triticale grass is a good source of antioxidants.

Compared with rye and triticale, wheatgrass and BG contained more amino acids, organic acids, nucleic acids, lipids and fatty acid in higher contents, demonstrating that wheatgrass and BG are better suited to develop functional foods.

Five compounds were detected in both positive and negative mode, L-isoleucine, malic acid, 9(S)-HOTrE, 9(S)-HpODE and 9(S)-HpOTrE, at high levels and are therefore important for grass growth and development. We also detected biochemicals unique to the individual grasses. In addition to the abovementioned metabolites, rye and triticale were rich in stearidonic acid, gallic acid, 9(S)-HpOTrE, ferulic acid and ketoleucine. Dehydroascorbic acid was only expressed in triticale in a high amount, levoglucosan and dodecanedioic acid were only expressed in rye in high amounts, and DIBOA-glucoside, MG(0:0/18:1(11Z)/0:0) and DIMBOA-glucoside were detected in wheat. This genetic variation could be the basis for improvements in cereal-based foods (Ziegler et al., 2016).

## Conclusions

LC-MS was applied to determine the untargeted metabolomic profiles of methanol extracts of grasses from four cereal species: wheat, barley, rye and triticale. We identified far more metabolites than previously reported and clear differences between the examined species. A total of 164 DEMs were detected, including amino acids, organic acids, lipids, fatty acids, nucleic acids, flavonoids, amines, polyamines, vitamins, sugar derivatives and other compounds, which revealed that the grasses contain diverse compositional makeups and nutrient distributions. Each species had unique metabolites in its grass. This study provides a glimpse into the metabolomics profiles of cereal grasses in China, which can form an important basis for nutrition, health and other parameters.

## Supplementary Files

**Supplemental Figure S1**. Total ion current chromatogram of a QC sample (positive mode)

**Supplemental Figure S2**. Total ion current chromatogram of a QC sample (negative mode)

**Supplemental Figure S3**. Total ion current chromatograms of HXH (positive mode)

**Supplemental Figure S4**. Total ion current chromatograms of HXH (negative mode)

**Supplemental Figure S5**. OPLS-PCA analysis between barley (BAR) and the three wheat cultivars (HXH, JM, and YN)

**Supplemental Figure S6**. OPLS-PCA analysis between rye (109) and the three wheat cultivars (HXH, JM, and YN).

**Supplemental Table S1**. DEM classification in positive mode

**Supplemental Table S2**. DEM classification in negative mode

## Acknowledgements

This work was supported by the Shandong Provincial Agricultural Elite Variety Project (2016LZGC010-3), National Natural Science Foundation of China (31671675) and Natural Science Foundation of Shandong Province under Grant (ZR2015CM034).

## Author Contributions

P. Xing and Z. Song designed the study and drafted the manuscript. P. Xing and X. Li collected test data and interpreted the results.

