## Supplemental Figure and tables for "Differences in the metabolite profiles of tender leaves of wheat, barley, rye and *Triticale* based on LC-MS"

SUPPLEMENTAL DATA

**Supplemental Figure S1** Total ion current chromatogram of QC sample (POS)

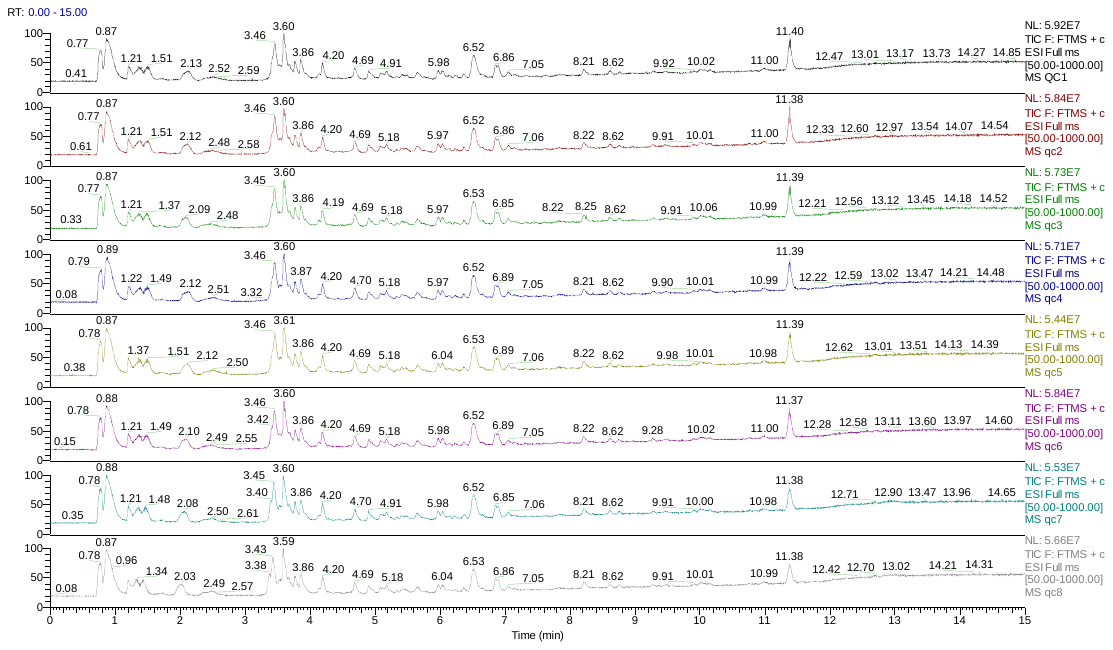

**Fig S1** Total ion current chromatogram of QC sample (POS)

**Supplemental Figure S2** Total ion current chromatogram of QC sample (NEG)

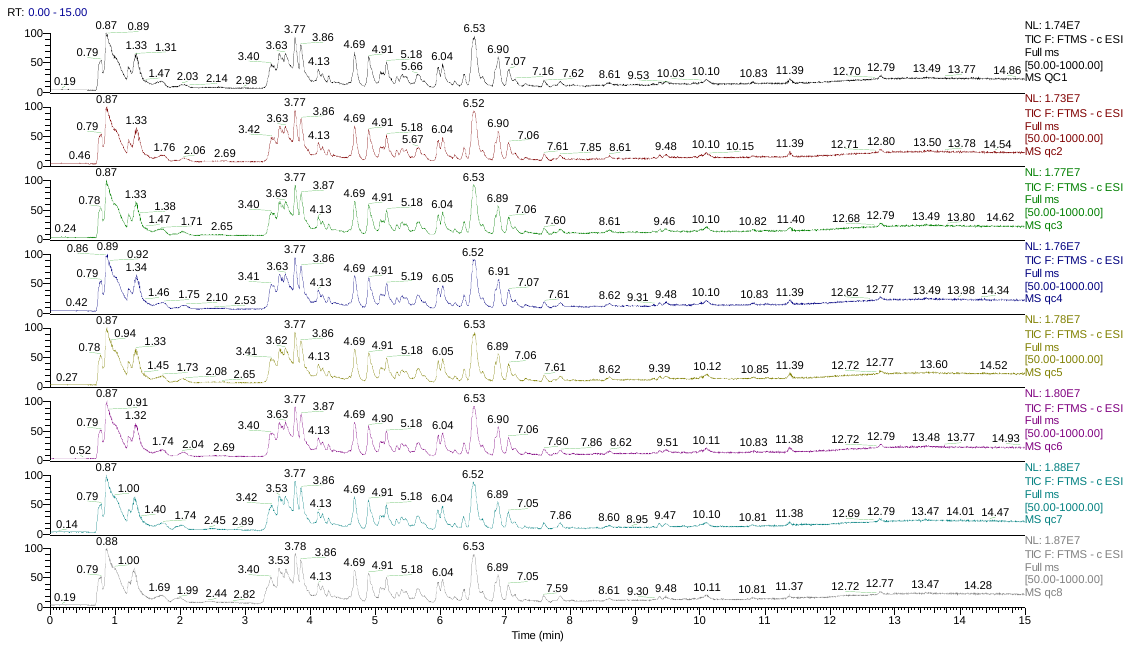

**Figure S2** Total ion current chromatogram of QC sample (NEG)

**Supplemental Figure S3** Total ion current chromatograms of metabolomic analysis for HXH in positive mode

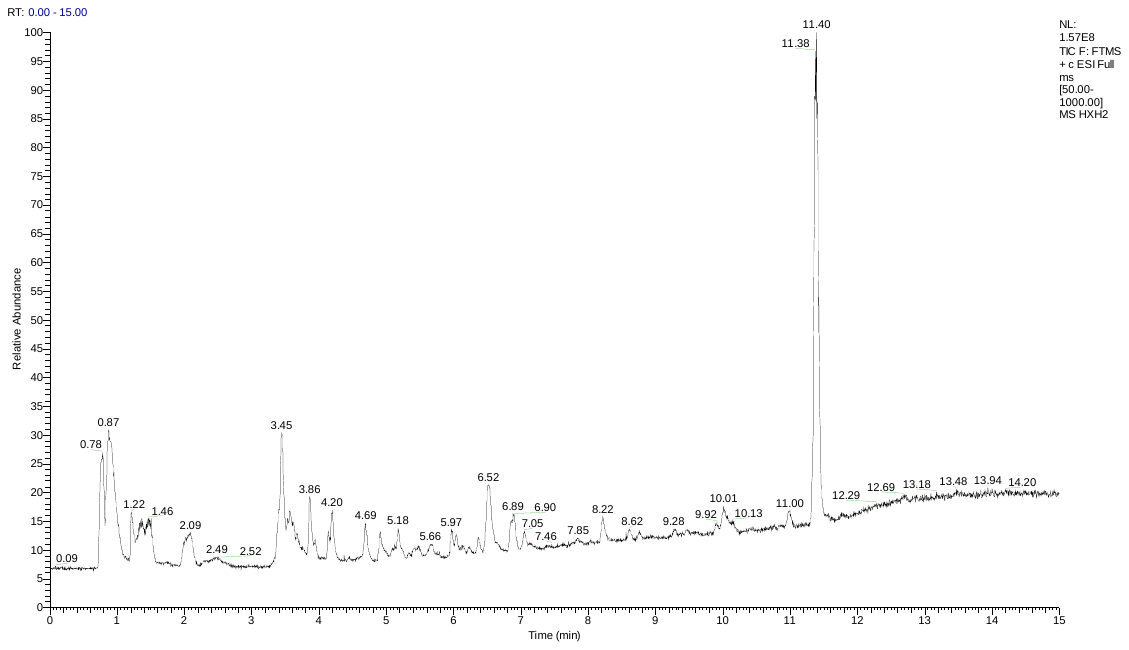

**Figure S3** Total ion current chromatograms of metabolomic analysis for HXH in positive scan mode

**Supplemental Figure S4** Total ion current chromatograms of metabolomic analysis for HXH in negative mode

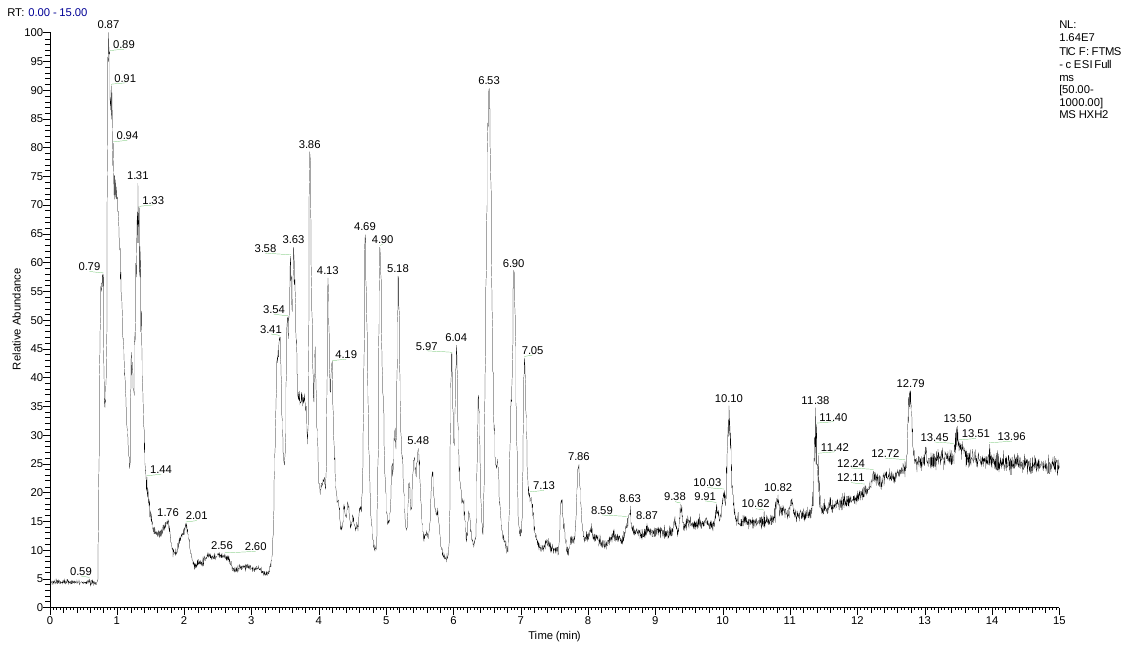

**Fig S4** Total ion current chromatograms of metabolomic analysis for HXH in negative scan mode (b)

**Supplemental Figure S5** OPLS-PCA analysis between barley (BAR) and three wheat cultivars (HXH, JM, YN)

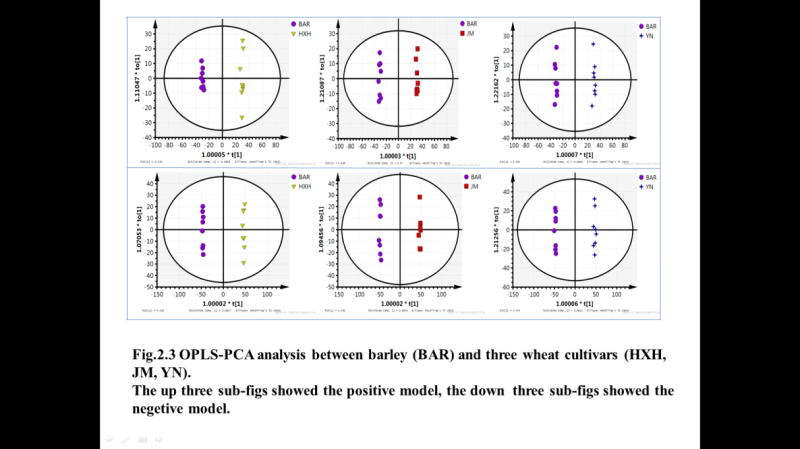

**Fig S5** OPLS-PCA analysis between barley (BAR) and three wheat cultivars (HXH, JM, YN).

Above three sub- figure showed the positive model, the below three sub-figures showed the negative model.

**Supplemental Fig S6** OPLS-PCA analysis between rye (109) and three wheat cultivars (HXH, JM, YN).

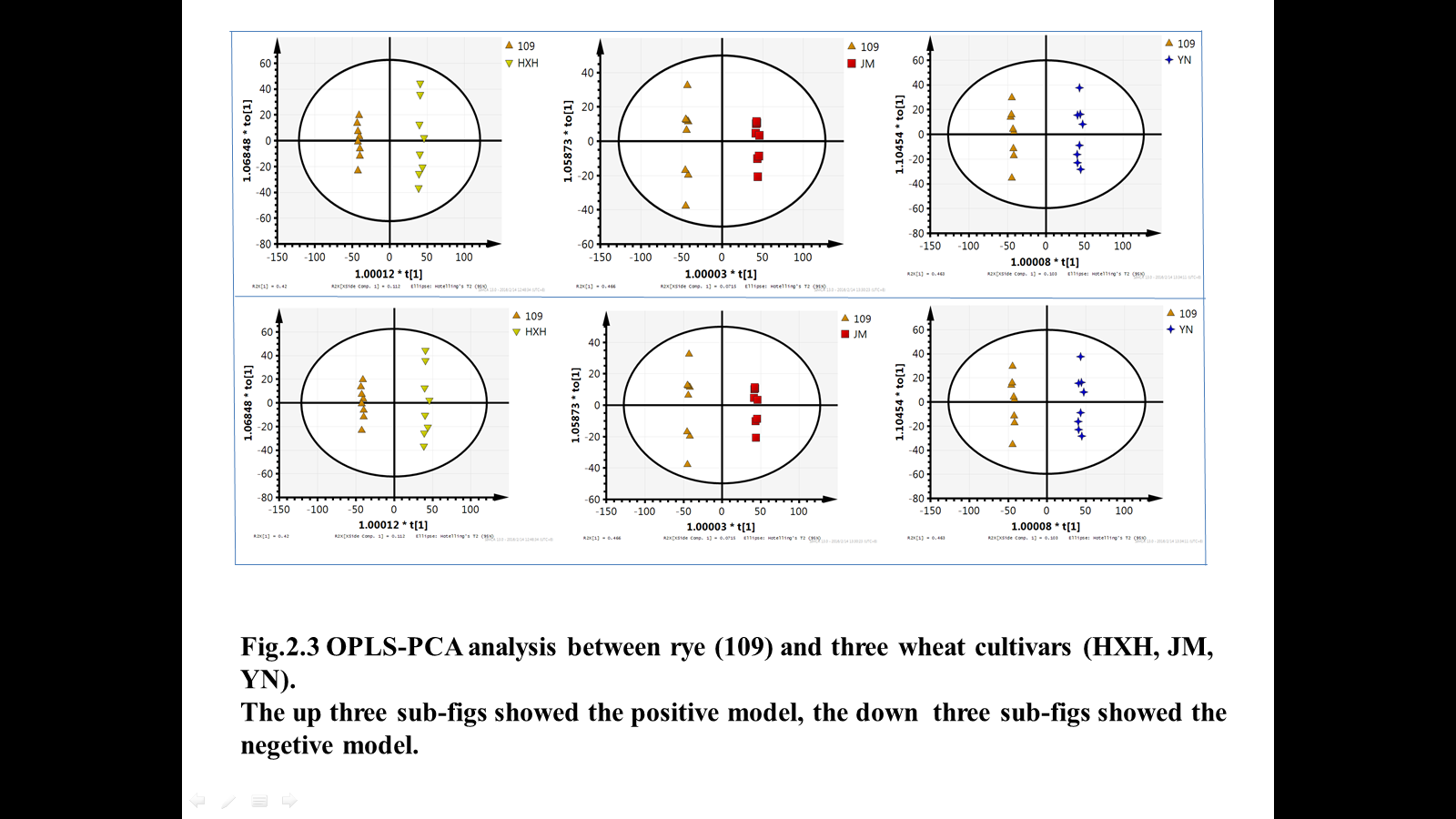

**Fig S6 OPLS-PCA analysis between rye (109) and three wheat cultivars (HXH, JM, YN).**

Above three sub- figure showed the positive model, the below three sub-figures showed the negative model.

**Supplemental Table S1**

**Supplemental Table S1 DEMs classification in positive mode**

| **Compound** | **Time** | **CompMW** | **Species with higher content** |
| --- | --- | --- | --- |
| **Amino acid** |  |  |  |
| L-Lysine | 0.805 | 146.11 | wheat |
| L-Tyrosine | 0.949 | 181.07 | wheat |
| L-Arginine | 0.841 | 174.11 | wheat |
| L-Isoleucine | 1.317 | 131.09 | wheat |
| Glutathione | 1.320 | 307.08 | wheat |
| L-Proline | 0.897 | 115.06 | Barley |
| L-Asparagine | 0.889 | 132.05 | Barley |
| L-Glutamine | 0.875 | 146.07 | Barley |
| L-Histidine | 0.809 | 155.07 | Barley |
| Ornithine | 4.013 | 132.09 | Barley |
| L-Theanine | 1.352 | 174.10 | Barley |
| 5-Hydroxy-L-tryptophan | 2.124 | 220.08 | Barley |
| L-Valine | 0.921 | 117.08 | Rye |
| L-Threonine | 0.877 | 119.06 | Rye |
| L-Glutamate | 0.893 | 147.05 | Rye |
| L-Methionine | 1.211 | 149.05 | Rye |
| L-Phenylalanine | 1.915 | 165.08 | Rye |
| L-Aspartic Acid | 0.895 | 133.04 | Triticale |
| β-Alanine | 0.865 | 89.05 |  |
| L-Serine | 0.872 | 105.04 |  |
| **Organic Acid** |  |  |  |
| Abscisic Acid | 1.752 | 264.13 | wheat |
| Nonanedioic acid | 4.133 | 188.10 | wheat |
| Deoxycholic acid | 3.570 | 392.28 | wheat |
| Phenylpyruvic acid | 3.896 | 164.05 | wheat |
| Caffeic Acid | 3.638 | 180.04 | wheat |
| DL-pipecolic acid | 0.900 | 129.08 | wheat |
| γ-Aminobutryic acid | 0.866 | 103.06 | Barley |
| α-ketoisovaleric acid | 3.430 | 116.05 | Barley |
| Pyroglutamic acid | 1.212 | 129.04 | Barley |
| Indoleacrylic acid | 3.305 | 187.06 | Barley |
| Jasmonic acid | 4.298 | 210.13 | Barley |
| Valerenic acid | 4.721 | 234.16 | Barley |
| ferulic acid | 3.559 | 194.06 | Rye |
| Gentisic acid | 3.559 | 154.03 | Rye |
| Vanillic acid | 3.748 | 168.04 | Triticale |
| Salicylic acid | 2.389 | 138.03 | Triticale |
| Malic acid | 1.259 | 134.02 | Triticale |
| Gallic acid | 1.937 | 170.02 | wheat and Triticale |
| p-Coumaroyl quinic acid | 3.577 | 338.10 | wheat and Triticale |
| Phenylacetic acid | 3.963 | 136.05 |  |
| **Llipids or fatty acid** |  |  |  |
| MG(0:0/18:1(11Z)/0:0) | 10.127 | 356.29 | wheat |
| MG(0:0/20:5(5Z,8Z,11Z,14Z,17Z)/0:0) | 9.455 | 376.26 | wheat |
| MG(0:0/20:4(5Z,8Z,11Z,14Z)/0:0) | 10.325 | 378.27 | wheat |
| MG(0:0/18:2(9Z,12Z)/0:0) | 7.109 | 354.28 | wheat |
| MG(16:0/0:0/0:0) | 10.088 | 330.28 | wheat |
| LysoPE(0:0/16:0) | 7.832 | 453.29 | wheat |
| LysoPC(16:0) | 8.058 | 495.33 | wheat |
| 9(S)-HODE | 8.441 | 296.23 | Barley |
| MG(18:0/0:0/0:0) | 11.354 | 358.31 | Barley |
| MG(0:0/20:3(5Z,8Z,11Z)/0:0) | 11.339 | 380.29 | Barley |
| DG(14:0/20:3(5Z,8Z,11Z)/0:0) | 14.192 | 590.49 | Barley |
| DG(18:3(6Z,9Z,12Z)/18:3(9Z,12Z,15Z)/0:0) | 13.484 | 612.47 | Barley |
| Stearidonic Acid | 4.907 | 276.21 | Barley |
| Homoserine lactone | 0.818 | 101.05 | Barley |
| α-Linolenic Acid | 7.604 | 278.22 | Rye |
| 9(S)-HOTrE | 6.906 | 294.22 | Rye |
| 9(S)-HpOTrE | 6.528 | 310.21 | Rye |
| 9(S)-HpODE | 7.054 | 312.23 | Rye |
| MG(0:0/18:3(6Z,9Z,12Z)/0:0) | 6.574 | 352.26 | Rye |
| MG(0:0/18:4(6Z,9Z,12Z,15Z)/0:0) | 9.665 | 350.24 |  |
| **Nucleic acid** |  |  |  |
| Dihydrouracil | 3.388 | 114.05 | Barley |
| Cytidine | 0.893 | 243.08 | Barley |
| Deoxyadenosine | 1.222 | 251.10 | Barley |
| Hypoxanthine | 3.786 | 136.04 | Rye |
| Xanthosine | 4.439 | 284.08 | Triticale |
| Thymidine | 1.763 | 242.09 | Triticale |
| Thymine | 1.679 | 126.04 | Triticale |
| 5-Methylcytosine | 1.210 | 125.06 | Triticale |
| Adenosine | 1.260 | 267.10 | Triticale |
| Cytosine | 1.207 | 111.04 | Triticale |
| Adenine | 1.159 | 135.05 |  |
| Uracil | 1.211 | 112.03 |  |
| Guanine | 0.914 | 151.05 |  |
| cGMP | 1.210 | 345.05 |  |
| **Vitamin** |  |  |  |
| D-Biotin | 2.422 | 244.09 | wheat |
| Pantothenic Acid | 2.362 | 219.11 | Barley |
| Niacin (Nicotinic acid) | 1.203 | 123.03 | Barley |
| Ubiquinone (Q2) | 5.312 | 318.18 | Barley |
| Ubiquinone-1 | 4.182 | 250.12 | Barley |
| Pyridoxal (Vitamin B6) | 3.611 | 167.06 | Rye |
| Riboflavin (Vitamin B2) | 3.632 | 376.14 | Rye |
| **Flavonoid** |  |  |  |
| Salicin | 1.303 | 286.11 | wheat |
| Caffeyl alcohol | 4.068 | 166.06 | wheat |
| Methyl jasmonate | 5.648 | 224.14 | Barley |
| Catechin | 0.941 | 290.08 | Barley |
| Coumarin | 3.323 | 146.04 | Rye |
| Kaempferol | 3.623 | 286.05 | Rye |
| Pheophorbide a | 10.157 | 592.27 | Triticale |
| Quercitrin | 3.656 | 448.10 | Triticale |
| Rutin | 3.586 | 610.15 | Triticale |
| 12-Ketodeoxycholic acid | 12.262 | 390.28 | Rye and Triticale |
| cis-Jasmone | 4.958 | 164.12 | Rye and Triticale |
| Stigmasterol | 9.828 | 412.37 |  |
| **Amines or Polyamine** |  |  |  |
| Linoleamide | 6.382 | 279.26 | wheat |
| Phytosphingosine | 6.564 | 317.29 | wheat and barley |
| Tryptamine | 3.573 | 160.10 | Barley |
| Serotonin | 1.914 | 176.09 | Barley |
| Choline | 0.933 | 103.10 | Barley |
| Sphinganine | 6.518 | 301.30 | Barley |
| Histamine | 0.789 | 111.08 | Rye |
| Niacinamide | 1.210 | 122.05 | Rye |
| Salicylamide | 2.496 | 137.05 | Triticale |
| **Sugar derivative** |  |  |  |
| L-Xylulose | 3.859 | 150.05 | wheat |
| Glucose 6-phosphate | 0.973 | 260.03 | Barley |
| Furfural | 3.440 | 96.02 | Barley |
| Glucosamine 1-phosphate | 4.626 | 259.05 | Triticale |
| **Other** |  |  |  |
| Indole | 3.353 | 117.06 | Barley |
| Hydroxyhydroquinone | 0.907 | 126.03 | Barley |
| Porphobilinogen | 2.195 | 226.10 | Barley |
| Pyrocatechol | 2.502 | 110.04 | Triticale |

**Supplemental Table S2 DEMs classification in negative mode**

| **Compounds** | **Time** | **CompMW** | **Species with higher content** |
| --- | --- | --- | --- |
| **Amino acid** |  |  |  |
| L-Threonine | 0.853 | 119.059 | wheat |
| L-Isoleucine | 1.431 | 131.095 | wheat |
| L-Lysine | 0.805 | 146.106 | wheat |
| L-Tyrosine | 1.214 | 181.074 | wheat |
| Ornithine | 0.822 | 132.090 | wheat |
| Pyroglutamic acid | 0.854 | 129.043 | wheat |
| L-Serine | 0.872 | 105.043 | wheat |
| L-Glutamate | 0.893 | 147.052 | wheat |
| L-Glutamine | 0.875 | 146.069 | wheat and barley |
| L-Histidine | 0.809 | 155.070 | wheat and barley |
| L-Asparagine | 0.889 | 132.054 | wheat and barley |
| L-Tryptophan | 3.759 | 204.090 | barley |
| β-Alanine | 1.254 | 89.048 | barley |
| N-Acetyl-L-glutamic acid | 1.212 | 189.064 | barley |
| 5-Hydroxy-L-tryptophan | 2.132 | 220.085 | barley |
| Ketoleucine | 3.733 | 130.063 | barley |
| L-Aspartic Acid | 0.895 | 133.040 | barley |
| L-Theanine | 1.347 | 174.101 | barley |
| Glutathione, oxidized | 0.918 | 612.151 | barley |
| L-Methionine | 1.210 | 149.051 | Rye |
| L-Valine | 3.418 | 117.080 | Rye and Triticale |
| L-Phenylalanine | 1.915 | 165.079 | Rye and Triticale |
| Acetyl-L-tyrosine | 3.419 | 223.084 | wheat and Triticale |
| **Organic Acid** |  |  |  |
| Cinnamic acid | 3.884 | 148.053 | wheat |
| β-Hydroxypyruvic acid | 1.160 | 104.011 | wheat |
| Nonanedioic acid | 4.133 | 188.105 | wheat |
| p-Coumaroyl quinic acid | 3.497 | 338.100 | wheat |
| Traumatic Acid | 4.292 | 228.136 | wheat |
| Vanillic acid | 3.584 | 168.042 | wheat |
| Gallic acid | 2.042 | 170.022 | wheat |
| Acetoacetic acid | 0.847 | 102.032 | wheat and Triticale |
| ferulic acid | 3.540 | 194.058 | wheat and Triticale |
| Chlorogenic Acid | 3.740 | 354.095 | wheat and Triticale |
| Salicylic acid | 2.389 | 138.032 | wheat and Triticale |
| Sinapic acid | 3.989 | 224.068 | wheat and Triticale |
| Octadecanedioic acid | 6.267 | 314.246 | barley |
| Methyl jasmonate | 5.646 | 224.141 | barley |
| Indoleacrylic acid | 3.305 | 187.063 | barley |
| Malic acid | 1.259 | 134.022 | barley |
| Pyruvate | 1.000 | 88.016 | barley |
| γ-Aminobutryic acid | 0.866 | 103.063 | barley |
| Abscisic Acid | 4.128 | 264.136 | barley |
| Jasmonic acid | 4.291 | 210.126 | barley |
| Dodecanedioic acid | 4.031 | 230.152 | barley |
| Succinic acid | 1.309 | 118.027 | barley |
| Fumaric acid | 1.241 | 116.011 | barley |
| α-ketoisovaleric acid | 3.430 | 116.047 | barley |
| Quinic acid | 0.888 | 192.064 | Triticale |
| Glutaconic acid | 1.307 | 130.027 | Triticale |
| Phenylpyruvic acid | 3.898 | 164.048 | Rye and Triticale |
| DL-α-Lipoic Acid | 3.668 | 206.043 | Rye and Triticale |
| Gentisic acid | 3.560 | 154.027 | Rye and Triticale |
| Glyoxylic acid | 3.787239552 | 74.001 | Rye and Triticale |
| Oxoglutaric acid | 1.202 | 146.022 |  |
| Phenylacetic acid | 3.963 | 136.053 |  |
| **Llipids or fatty acid** |  |  |  |
| cis-9-palmitoleic acid | 9.474 | 254.224 | wheat |
| Palmitic acid | 8.575 | 256.240 | wheat |
| LysoPE(0:0/16:0) | 7.827 | 453.285 | wheat |
| LysoPE(0:0/18:3(6Z,9Z,12Z)) | 4.606 | 475.270 | wheat |
| LysoPE(0:0/18:2(9Z,12Z)) | 7.227 | 477.286 | wheat |
| LysoPE(0:0/18:1(11Z)) | 8.096 | 479.301 | wheat |
| LysoPC(15:0) | 8.049 | 481.317 | wheat |
| LysoPE(0:0/20:2(11Z,14Z)) | 8.371 | 505.316 | wheat |
| Gluconolactone | 2.426 | 178.041 | wheat |
| MG(0:0/15:0/0:0) | 7.072 | 316.261 | wheat |
| Oleic Acid | 11.026 | 282.256 | wheat and Triticale |
| α-Linolenic Acid | 9.652 | 278.224 | wheat and Triticale |
| MG(0:0/18:1(11Z)/0:0) | 10.127 | 356.292 | wheat and Triticale |
| Linoleic acid | 10.090 | 280.240 | wheat and Triticale |
| LysoPE(0:0/20:1(11Z)) | 9.157 | 507.332 | wheat and Triticale |
| MG(0:0/22:5(4Z,7Z,10Z,13Z,16Z)/0:0) | 9.564 | 404.293 | wheat and Triticale |
| 9(S)-HpODE | 7.052 | 312.230 | Rye |
| 9(S)-HOTrE | 8.298 | 294.219 | Rye |
| 9(S)-HODE | 8.951 | 296.235 | Rye |
| 9(S)-HpOTrE | 6.593 | 310.214 | Rye |
| Stearidonic Acid | 0.928 | 276.209 | Rye |
| **Nucleic acid** |  |  |  |
| Uracil | 1.207 | 112.027 | wheat |
| Hydrouracil | 0.850 | 128.059 | wheat |
| Deoxyinosine | 3.768 | 252.085 | wheat |
| cAMP | 1.209 | 329.052 | wheat |
| cGMP | 1.210 | 345.047 | wheat |
| Dihydrouracil | 3.388 | 114.043 | wheat and barley |
| Adenosine | 0.916 | 267.097 | wheat and Triticale |
| Thymidine | 1.763 | 242.090 | wheat and Triticale |
| 2'-Deoxyuridine | 1.240 | 228.075 | wheat and Triticale |
| dGMP | 4.103 | 347.064 | Triticale |
| Adenine | 1.177 | 135.055 |  |
| Xanthine | 1.212 | 152.034 |  |
| **Vitamin** |  |  |  |
| Niacin (Nicotinic acid) | 3.864 | 123.032 | wheat |
| Pantothenic Acid | 2.362 | 219.111 | barley |
| Ubiquinone-1 | 4.298 | 250.118 | barley |
| Dehydroascorbic acid | 0.990 | 174.017 | Triticale |
| Pyridoxal (Vitamin B6) | 3.866 | 167.058 | Triticale |
| 4-Pyridoxic acid | 2.789 | 183.053 | Rye |
| Riboflavin (Vitamin B2) | 3.632 | 376.136 |  |
| flavonoid |  |  |  |
| Catechin | 1.232 | 290.075 | wheat |
| Quercitrin | 3.995 | 448.101 | wheat |
| cis-Jasmone | 4.524 | 164.120 | barley |
| Caffeyl alcohol | 4.005 | 166.063 | barley |
| Rutin | 3.593 | 610.153 | Triticale |
| Pheophorbide a | 10.156 | 592.268 | Triticale |
| Sinapyl alcohol | 4.132 | 210.091 | Triticale |
| Kaempferol | 3.475 | 286.048 | Rye and Triticale |
| Baicalin | 3.936 | 446.085 | Rye |
| **Sugar derivative** |  |  |  |
| Sedoheptulose 7-phosphate | 0.934 | 290.041 | wheat and Triticale |
| L-Xylulose | 3.859 | 150.053 | wheat and Triticale |
| D-Mannitol | 0.844 | 182.079 | wheat |
| Quercetin 3-O-glucoside | 4.241 | 464.095 | wheat |
| Levoglucosan | 0.892 | 162.053 | barley |
| Glucoheptonic acid | 0.845 | 226.069 | Triticale |
| DIMBOA-glucoside | 3.652 | 373.100 | Triticale |
| Gluconic acid | 0.881 | 196.059 | Rye |
| DIBOA-glucoside | 3.432 | 343.098 | Rye and Triticale |
| 6-Acetyl-D-glucose | 3.394 | 222.074 |  |
| **Amines or polyamine** |  |  |  |
| Histamine | 0.789 | 111.080 | wheat and barley |
| Phosphorylcholine | 3.797 | 169.051 | wheat and barley |
| Nicotianamine | 0.906 | 303.143 | Rye |
| Salicylamide | 2.496 | 137.041 | Rye and Triticale |
| **Other** |  |  |  |
| Hydroxyhydroquinone | 3.262 | 126.032 | wheat |
| L-Dopa | 3.704 | 197.065 | wheat |
| Indole | 3.945 | 117.058 | barley |
| Porphobilinogen | 2.174 | 226.095 | barley |
| Furfural | 3.440 | 96.021 | Triticale |
| Pyrocatechol | 3.561 | 110.037 | Rye and Triticale |
